# Social learning of navigational routes in tandem running acorn ants (*Temnothorax nylanderi*)

**DOI:** 10.1101/2024.06.05.597530

**Authors:** Aina Colomer-Vilaplana, Tara Williams, Simone M. Glaser, Christoph Grüter

**Affiliations:** Institute of Organismic and Molecular Evolutionary Biology, Johannes-Gutenberg University of Mainz, Germany; Department of Genetics and Microbiology, Universitat Autònoma de Barcelona, Bellaterra, Barcelona, 08193, Spain; Institute of Biotechnology and Biomedicine, Universitat Autònoma de Barcelona, Bellaterra, Barcelona, 08193, Spain; School of Biological Sciences, University of Bristol, 24 Tyndall Avenue, BS8 1TQ Bristol, UK

**Keywords:** ant, recruitment, navigation, tandem running, social learning, *Temnothorax nylanderi*

## Abstract

Tandem running in ants is a form of social learning that involves an informed leader guiding a naïve nestmate to a valuable resource, such as a nest site or a food source. Little is currently known about what tandem followers learn and how socially acquired navigational information affects future trips. While some studies suggest that tandem followers learn the resource position but not the route taken by the tandem pair to reach the resource, more recent evidence contradicts this view. We studied tandem running in foraging acorn ants *Temnothorax nylanderi* and provide evidence that tandem followers socially learn routes from their leaders and later use these routes when travelling between their nest and a food source. Followers that became tandem leaders themselves then guided their follower along the same routes in 90% of tandem runs, demonstrating that navigational information can spread in a forager population through sequential social learning. Ants increased their travelling speed, but not path straightness over successive trips. We also found that ants needed less time on subsequent trips if they experienced longer-lasting tandem runs, suggesting that longer lasting tandem runs allow followers to learn routes more efficiently. Adding visual cues did not affect most of the quantified variables, and we currently know little about the cues used by *T. nylanderi* during navigation. We discuss how the visual environment inhabited by different species might affect the importance of route learning during tandem running.

## Introduction

Social learning shapes the behaviour of animals in a wide range of biological contexts as it allows animals to acquire behaviours that boost survival and reproduction (Heyes, 2012; Hoppitt & Laland, 2013; Kendal et al., 2018; Laland, 2004). Social insects frequently rely on social information, provided in the form of signals or incidental cues, for example when foraging, during colony migrations and for nest defence (Grüter & Leadbeater, 2014; Leadbeater & Chittka, 2007; Leadbeater & Dawson, 2017).

A relatively well-studied example of social learning in social insects is tandem running in ants: after an ant has discovered a good food source or a suitable nest site, she returns to her nest to guide a fellow nestmate to the discovered resource (Franks & Richardson, 2006; Hingston, 1929; Kaur et al., 2012; Möglich et al., 1974; Silva et al., 2021; Wilson, 1959; reviewed by Franklin, 2014; Sasaki & Pratt, 2021). During foraging, tandem running may help colonies visit food sources of better quality and defend them as a group against competitors (Glaser et al., 2021; Glaser & Grüter, 2023; Goy et al., 2021; Shaffer et al., 2013). During nest relocations, tandem running helps colonies assess the quality of new nest sites and migrate more efficiently (Mizumoto et al., 2023; Stuttard et al., 2015). Tandem running has been described in dozens of ant species and is likely to have evolved several times independently in phylogenetically distant ant groups belonging to at least 4 subfamilies (Glaser & Grüter, 2022; Mizumoto et al., 2023). A common feature of ant species that use tandem running is that they have relatively small colony sizes (Beckers et al., 1989; Glaser & Grüter, 2022).

To initiate a tandem run inside the nest, tandem leaders of some (and maybe all) species produce a short-range pheromone that alerts potential followers to the discovery of a resource and attracts them to the tandem leader (“tandem calling”; Möglich et al., 1974). Ant pairs then walk towards their goal, with the follower ant frequently touching the legs and abdomen of the leader to signal her presence, while the leader continues to release pheromone to help followers maintain contact with the leader (Basari, Laird-Hopkins, et al., 2014; Franks et al., 2022; Hölldobler & Traniello, 1980; Möglich et al., 1974; Stuttard et al., 2015). Because tandem leaders invest time to help a nestmate learn the location of a resource, tandem running is often considered a case of animal teaching (Franks & Richardson, 2006; Richardson et al., 2007). It is still unclear, however, what ants learn when they follow a tandem leader and how this information affects future navigation. Followers might learn the *target location in relation to the nest location* and return to their nest using path-integration or environmental cues, such as landmarks or local odours. Ants can perform path integration by using a celestial compass, to gauge direction, in combination with a step-counting mechanism to estimate distances (Wittlinger et al., 2006). This type of navigational learning has been studied extensively in desert ants (esp. *Cataglyphis* and *Melophorus*, reviewed in Collett et al., 2013; Knaden & Graham, 2016; Zeil, 2012). Alternatively, tandem followers could learn *the route* itself and later attempt to retrace their steps when returning home, relying on environmental or idiothetic (self-generated) cues (Knaden & Graham, 2016; Sasaki et al., 2020). It has been argued that tandem followers do not learn routes in rock ants (*Temnothorax albipennis*), the only ant species where this has been studied in more detail, as ants did not return to their nest using a similar path to the one taken by the tandem pair (Basari, Bruendl, et al., 2014; Franklin, 2014; Franklin & Franks, 2012; Franks & Richardson, 2006). Instead, these studies suggested that *T. albipennis* tandem followers learn target locations and subsequently use path integration and visual cues for homing using different routes. In contrast, Sasaki et al. (2020) recently provided the first quantitative analysis of paths of former tandem followers and found that tandem followers did learn routes from their leaders, which they subsequently used when walking from their nest to the food source, but not when they returned from the food source to the nest. The finding that the learning of specific routes was only apparent when ants travelled towards the food, but not towards the nest is consistent with the earlier observations that ants returning to their nest did not use the tandem route (Basari, Bruendl, et al., 2014; Franklin & Franks, 2012; Franks & Richardson, 2006). Visual cues seem to play an important role in *T. albipennis* ants navigating between nest and resource (McLeman et al., 2002; Pratt et al., 2001).

It is currently unknown if learning during tandem following is similar in other ant species. Here, we assessed learning during tandem following in the acorn ant *Temnothorax nylanderi*, a species inhabiting woodland habitats (Seifert, 2018), using two different experimental set-ups. First, we tested ants in a binary choice setup up that offered ants two alternative routes to a food source, mimicking a situation where the most direct path from nest to food is blocked by an obstacle. In the second setup, ants navigated in an open area, *i.e.* an environment without obstacles. Since visual cues have been shown to affect navigation in other *Temnothorax* species (*T. unifasciatus*: Aron et al. 1988; *T. albipennis*: Pratt et al. 2001; McLeman et al. 2002; *T. rugatulus*: Bowens et al. 2013) and the tandem running *Diacamma indicum* (Mukhopadhyay & Annagiri, 2021), we tested ants in visually enriched and non-enriched environments. In addition, we also tested if former followers would teach routes they learned to naïve followers in later trips.

## Methods

### Study species and maintenance

Twenty-six colonies were collected from small decaying branches in the Lenneberg forest near Mainz, Germany (Experiment 1: 13 colonies collected between September 2020 and December 2021; colony size = 121 ± 20.0 (mean ± SD); Experiment 2: 13 colonies were collected between November 2014 and April 2016; colony size = 125 ± 28.3). All colonies had a reproductive queen and brood. Ants were kept in artificial nests made of two microscope slides (50mm×10mm×3mm) with a plexiglass slide in between the slides containing an oval living space and a nest entrance. Nests were covered with transparent red filter paper to reduce the light entering the living space. Each nest was stored in a nest box (100mm×100mm×30mm). Colonies were fed twice a week with a drop of honey and fruit flies or crickets. Water was available *ad libitum* throughout the study in a 1.5ml Eppendorf tube. Colonies were kept in a climate chamber at a temperature of 22°C, 70% humidity and a 12/12h light/dark cycle. Prior to a trial, colonies were starved of honey for 10 days to make sure colonies were motivated to forage.

#### Experiment 1

Focal colonies were placed in an experimental arena (20cm×30cm×1cm; cleaned with ethanol before adding a colony) (Fig. 1a) on day 7 of the starvation period to allow colonies to acclimatise to the experimental arena and chemically mark the territory (Bowens et al., 2013). The experimental arena presented a binary choice: ants could travel to a food source via a left or a right branch. The arena walls were covered with paraffin oil to prevent the ants from escaping. Each colony was tested once in a visually enriched environment (blue and yellow paper on arena floor) and once without visual enrichment (white paper placed beneath the arena; however, lab equipment and furniture in the laboratory provided visual landmarks) (Fig. 1a). Treatment order was randomised for each colony. Due to a malfunction of the climate cabinet, some colonies were damaged and only 21 of 26 possible trials could be performed.

**Figure 1:**
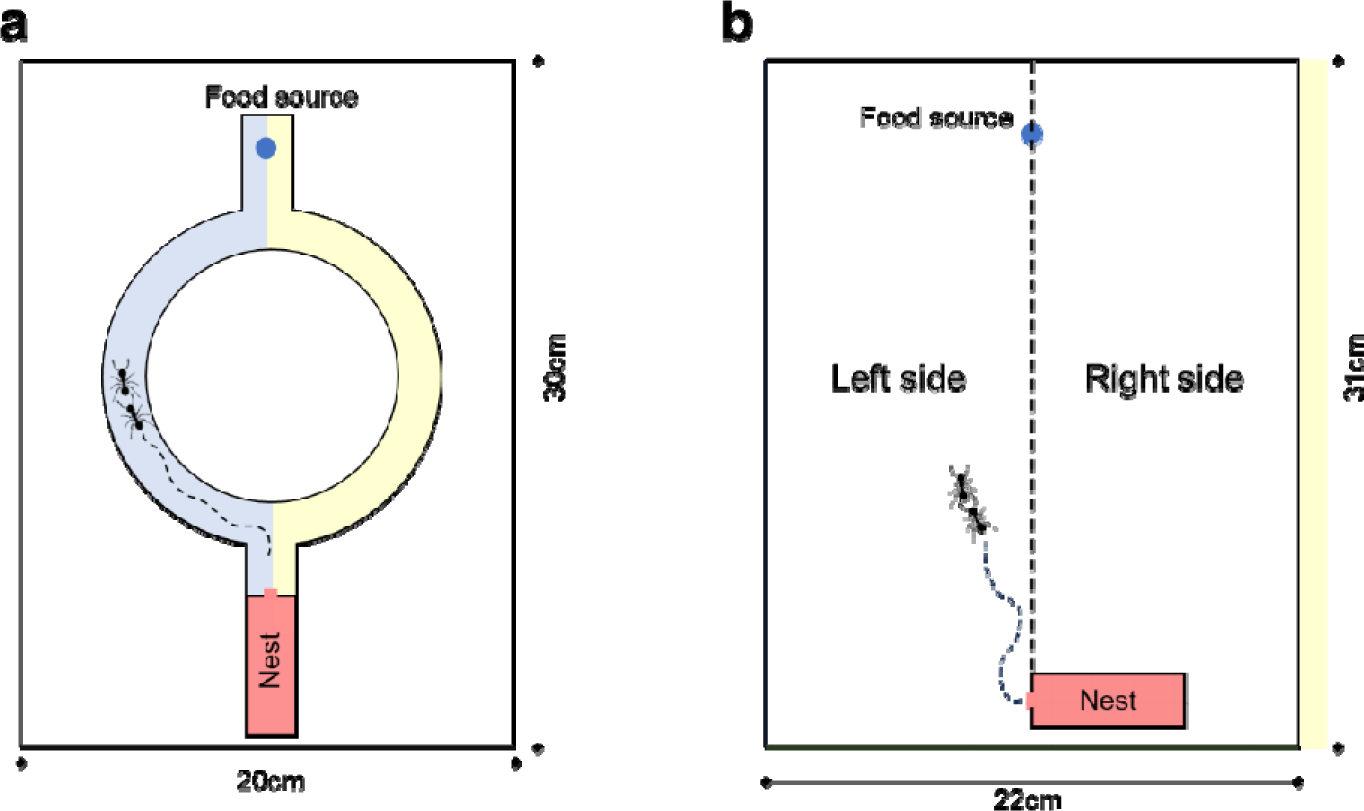
Experimental arenas used for Experiment 1 (a) and Experiment 2 (b) (a) Ants could use two branches to reach the food source. (b) Ants navigated an open foraging arena. A 1M sucrose solution was used as a food source. Each experiment tested two conditions, with and without added visual cues. Visual cues consisted of covering the floor with blue and yellow paper (a) or covering the sides of the arena (5cm height) with blue, green, black and yellow paper (b).

On a test day, a drop of unscented 1M sucrose solution was placed at the opposite end of the nest (25 cm from the nest; *T. nylanderi* typically forage less than 50 cm from their nest, see Heinze et al., 1996). A trial started when the first scout ant discovered the sucrose solution and lasted 120 minutes. We recorded trials with a HC-V130 Panasonic camera placed above the arena on a tripod. Followers of the first tandem runs (up to 10 followers per trial) were individually marked on the abdomen with different colour dots (Posca, Mitsubishi Pencil Co., UK) to allow individual ant identification throughout the trial. After a trial, colonies were returned to their nest boxes, provided with food and allowed a minimum of seven days recovery before being starved again.

From the video recordings, we recorded the branch choices (left or right) and trip durations of the tandem pairs and of follower’s subsequent eight journeys to the nest and the food source. Trip duration was measured as the time between leaving (or entering, in the case of a homing trip) the nest entrance and touching (or leaving, in the case of a homing trip) the sugar solution. We also recorded if former followers became leaders of tandem runs during a trial and we analysed the branch choice of these secondary tandem runs.

#### Experiment 2

Focal colonies were placed in on open experimental arena on day 4 of the starvation period to allow colonies to explore and get used to their new surroundings. The experimental arena consisted of a 31cm×22cm×5cm transparent box (cleaned with ethanol before adding a colony), with the walls partially covered with Fluon to prevent the ants from escaping (Fig. 1b).

Nests were placed on one side of the experimental arena, 25 cm from the food source location (Fig. 1b). A drop of 1M sucrose solution was again used as a food source. A trial started when the first scout ant discovered the sucrose solution and lasted 90 minutes. The experimental arena was recorded from above using a HC-V130 Full HD Panasonic camera. Tandem followers were individually marked as described above. After a trial, colonies were returned to their nest box, fed and allowed to recover as described above. Each colony was tested once with added visual cues (blue, green, black and yellow paper covering the arena walls) and once without added cues (white paper covering the arena walls). Treatment order was randomised for each colony.

We analysed the tandem runs of individually marked followers, their first return trip to the nest (trip 1) and the first return trip to the food source (trip 2). We focused on the first two trips because we assumed that these trips are most likely to be affected by experience gained during the tandem run. We measured trip duration, trip distance and speed (or rate of progress; cm/sec) using the object detection and tracking software *AnTracks* (www.antracks.org). To measure speed, we divided the total length of a trip (cm) by the duration (sec) of the trip. Fig. S1 shows the combined trajectories of all studied ants during a 90-minute trial.

To assess whether homing ants choose a route passing through the same areas in the arena as when they were tandem followers, we divided the arena into a left and right side (Fig. 1b). We measured the time ants spent on either side, arbitrarily dividing the time spent on the left side by the total trip time to calculate the percentage of time spent on the left side. This allowed us to test if the proportion of time spent on one side during the tandem run predicts the probability that an ant will walk on the same side when returning to the nest or the food source.

### Statistical analysis

#### Experiment 1

Statistical analyses were carried out in R 4.1.3 (R Core Team, 2021). To test if former followers used the same path on subsequent trips as during the tandem run, generalised linear mixed-effect models (GLMMs) with a binomial response (1 = same route as tandem run, 0 = different from tandem run) were used (lme4 package; Bates et al., 2015). Linear mixed-effects models (LMEs) with a normally distributed response variables were used to analyse trip durations. Visual condition (enriched vs. non-enriched) and trip number were included as fixed effects to explore their effects on branch choice and trip duration. We tested the significance of interactions between fixed effects using likelihood ratio tests (LRT) and removed non-significant interactions (Zuur et al., 2009). Colony ID and Ant ID were included as random effects to control for the non-independence of data points from the same ants and colonies. We checked our LMEs for normality and homogeneity of variance using the DHARMa package (Hartig, 2022) and used the box-cox method if necessary to optimally transform data (Crawley, 2007).

#### Experiment 2

We used GLMMs and LMEs to analyse the effects of visual condition (enriched vs. non-enriched) and tandem characteristics (duration, distance, speed and arena side) on the duration, distance, speed and arena side of subsequent trips (trip 1 and 2). Binomial GLMMs were used to test if the probability that ants spend the majority of time (>50%) on one side during trip 1 and 2 depended on time spent on the same side during the original tandem run. Response variables in LMEs were transformed using the box-cox transformation if necessary to comply with assumptions, as described above. We tested the significance of interactions between fixed effects as described above. When significant predictors had three levels, pairwise comparisons were performed using the multcomp package (Hothorn et al., 2008). We used the “Tukey” method to adjusts for multiple comparisons by controlling the family-wise error rate. Colony ID and Ant ID were again used as random effects.

### Ethical Note

No licences or permits were required for this research. During experiments the ants were not harmed and were taken care of as outlined in the *Study species and maintenance* section.

## Results

### Experiment 1

Overall, 125 tandem run followers were individually marked and observed. These tandem followers undertook a total of 599 trips. Tandem pairs had no significant preference for the left or right branch (GLMM, z-value = 0.77, p = 0.44). For the following analysis, we focused on the first eight trips after the initial tandem run (four return to nest trips, four return to food trips) because only very few ants performed more than eight trips.

For all trips, except trip 7 (nest return trip 4), there was a strong, significant preference to choose the same branch as the one experienced during the tandem run (Table 1; Fig. 2) and even when ants returned to the food source the fourth time (trip 8), they had >80% probability to choose the same branch as the one taken during the tandem run.

**Figure 2:**
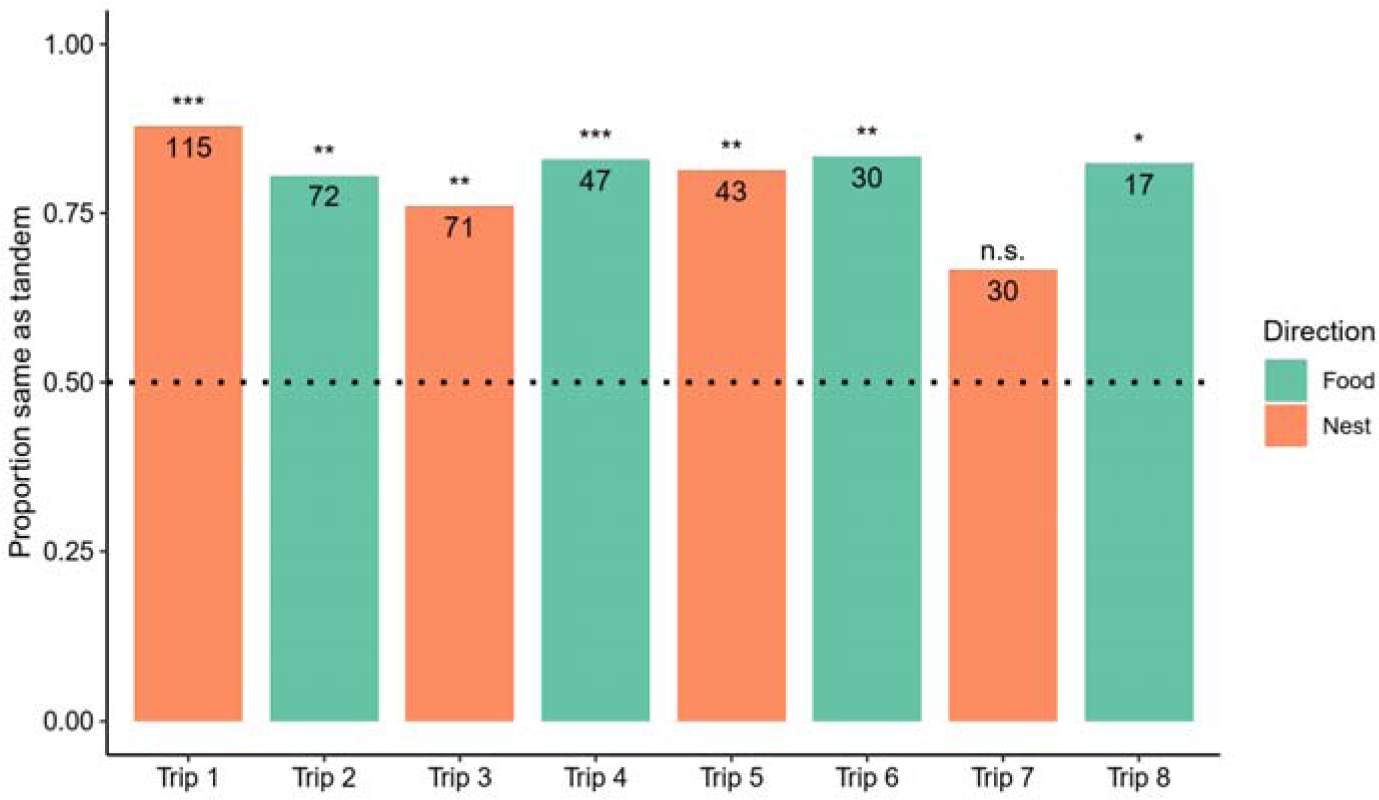
Probability that ants chose the same branch as the one taken when they were followers of a tandem run. Numbers indicate sample sizes, n.s. = not significant, ***p<0.0001, **p<0.01, *p<0.05.

**Table 1.** Trips performed by former followers. Percentage of ants choosing the same branch as experienced during that tandem run is shown.

| Trip | N | Same as tandem | z-value | p-value |
| --- | --- | --- | --- | --- |
| Trip 1 (nest return 1) | 115 | 87.8% | 6.93 | <0.0001 |
| Trip 2 (food return 1) | 72 | 80.6% | 3.77 | 0.0002 |
| Trip 3 (nest return 2) | 71 | 76.1% | 2.62 | 0.0088 |
| Trip 4 (food return 2) | 47 | 83.0% | 4.08 | <0.0001 |
| Trip 5 (nest return 3) | 43 | 81.4% | 2.61 | 0.009 |
| Trip 6 (food return 3) | 30 | 83.3% | 3.21 | 0.0013 |
| Trip 7 (nest return 4) | 30 | 66.7% | 1.79 | 0.07 |
| Trip 8 (food return 4) | 17 | 82.4% | 2.38 | 0.017 |

As trip number increased, the probability to choose the same branch as during their tandem run decreased slightly but significantly (Fig. 2) (GLMM: z-value = −2.59, p = 0.01). Visual enrichment had no significant effect on the probability to choose the same branch as during the tandem run (GLMM: z-value = - 0.18, p = 0.86; interaction: LRT = 0.39, p = 0.53).

We then tested if trip duration depended on trip number, travel direction and visual enrichment and found a significant interaction between trip number and travel direction (LME: LRT = 30.3, p <0.0001). Fig. 3 reveals that trips to the nest were faster than trips to the food source, except the first return to the nest (trip 1), which explains the significant interaction. Therefore, we analysed trip durations in both directions separately.

**Figure 3:**
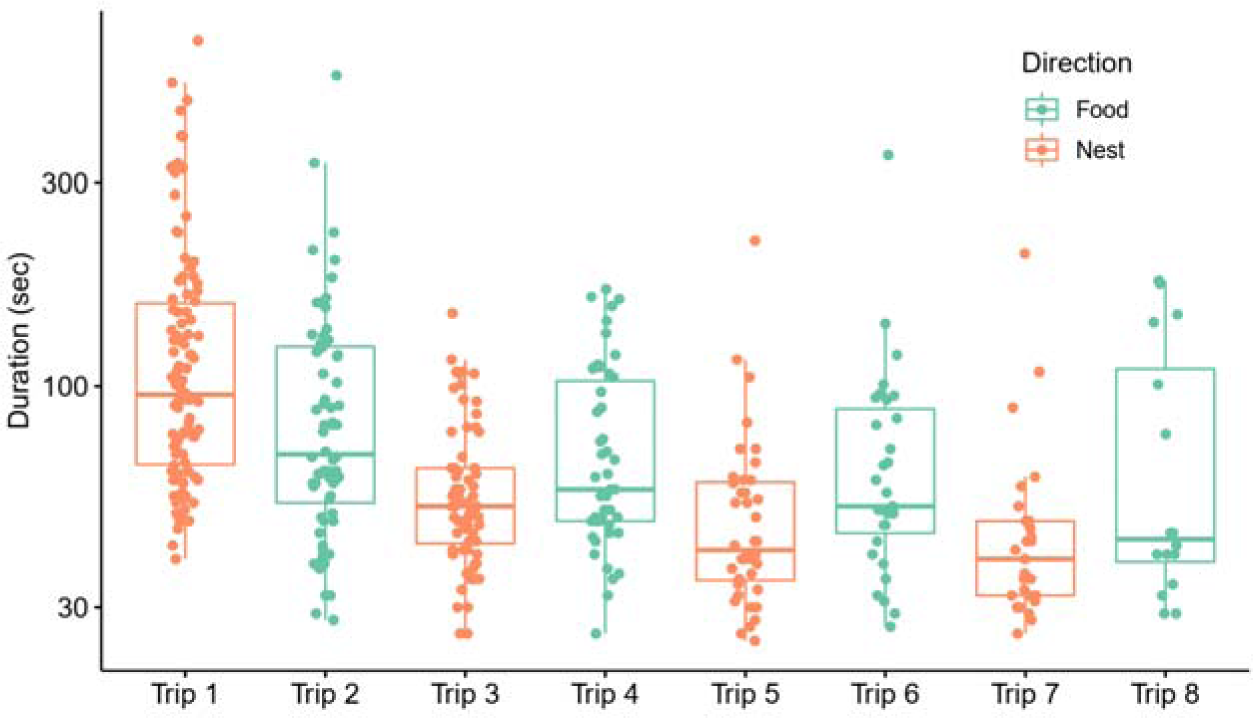
Trip duration in seconds for different trips and travel directions. Note that the y-axis is in log10-scale. Boxplots show median, 25% and 75% quartile and 5% and 95% percentile. Dots represent different ants.

Trips to the nest decreased in duration with increasing trip number (GLMM: t-value = 13.81, p < 0.0001), but there was no effect of added visual cues (t-value = 0.92, p = 0.36). Similarly, trips to the food became shorter over successive trips (t-value = 2.99, p = 0.003), but no effect of visual cue enrichment was found on trip duration to food (t-value = 0.54, p = 0.59).

Of all observed followers, 14 went on to lead tandem runs themselves. They led 30 tandem runs, and they were significantly more likely to use the same branch as the one they had experienced when they were tandem followers (90% choosing same branch, GLMM: z-value = 2.29, p = 0.022).

### Experiment 2

In the second experiment, ants navigated in an open arena, with or without the presence of additional visual cues on the walls of the arena. Overall, we observed 248 individually marked ants, 200 were tandem run followers and 46 were scouts that initially discovered the food source. Scouts were also studied to compare the homing time of social and individual learners (tandem followers vs. scouts).

#### Trip durations

We first compared the homing efficiency of ants that found the food source either by themselves (scouts; individual learners) or by following a tandem run (recruits, social learners) and found that former followers needed significantly less time than scouts (−20.2%; Fig. 4) (121.5 ± 54.6 sec vs. 97.0 ± 51.2 sec) (LME: t-value = 3.24, p = 0.0014; visual cues: t-value = −1.25, p = 0.21; interaction: LRT = 0.96, p = 0.33) to return to the nest.

**Figure 4:**
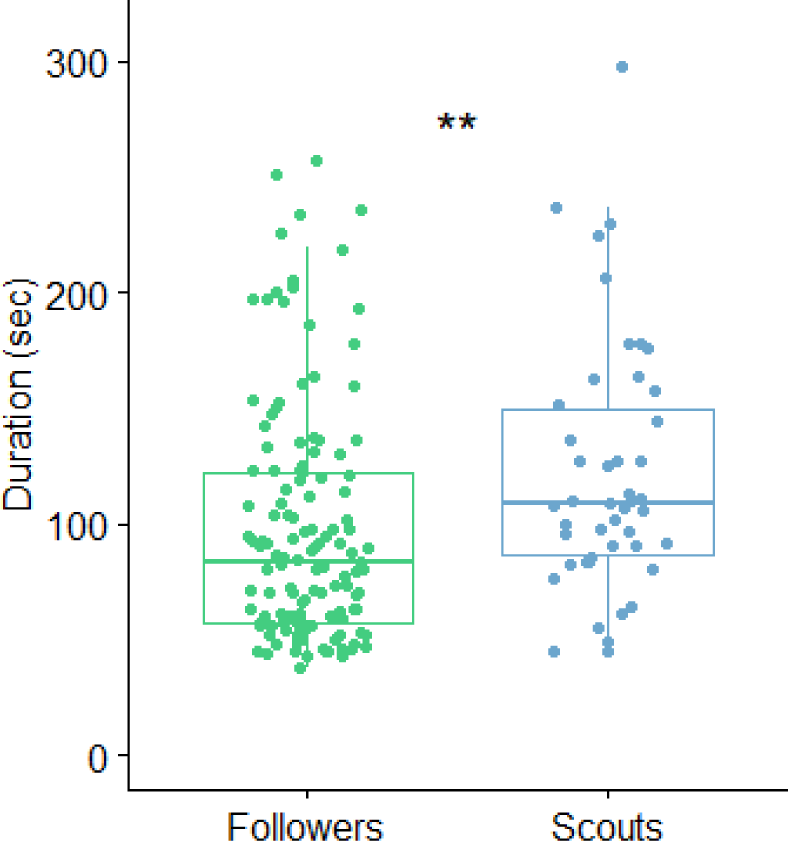
Duration of trips back to the nest (trip 1) of tandem followers and scouts.

We then analysed the characteristics of trips over the course of a trial and found that trip durations decreased (Fig. 5a) (LME: t-value = 17.65, p < 0.0001). Surprisingly, trips took longer when added visual cues were present (t-value = 4.75, p < 0.0001, interaction: LRT = 0.9, p = 0.34). The distance walked by ants did not change over time (Fig. 5b) (t-value = 1.62, p = 0.11), but trip distances were longer when visual cues were present (t-value = 5.21, p < 0.0001; interaction: LRT = 0.42, p = 0.51). The walking speed increased during the trial (Fig. 5c) (t-value = 27.34, p < 0.0001), independently on the presence of additional visual cues (t-value = 0.85, p = 0.40; interaction: LRT = 0.29, p = 0.59).

**Figure 5:**
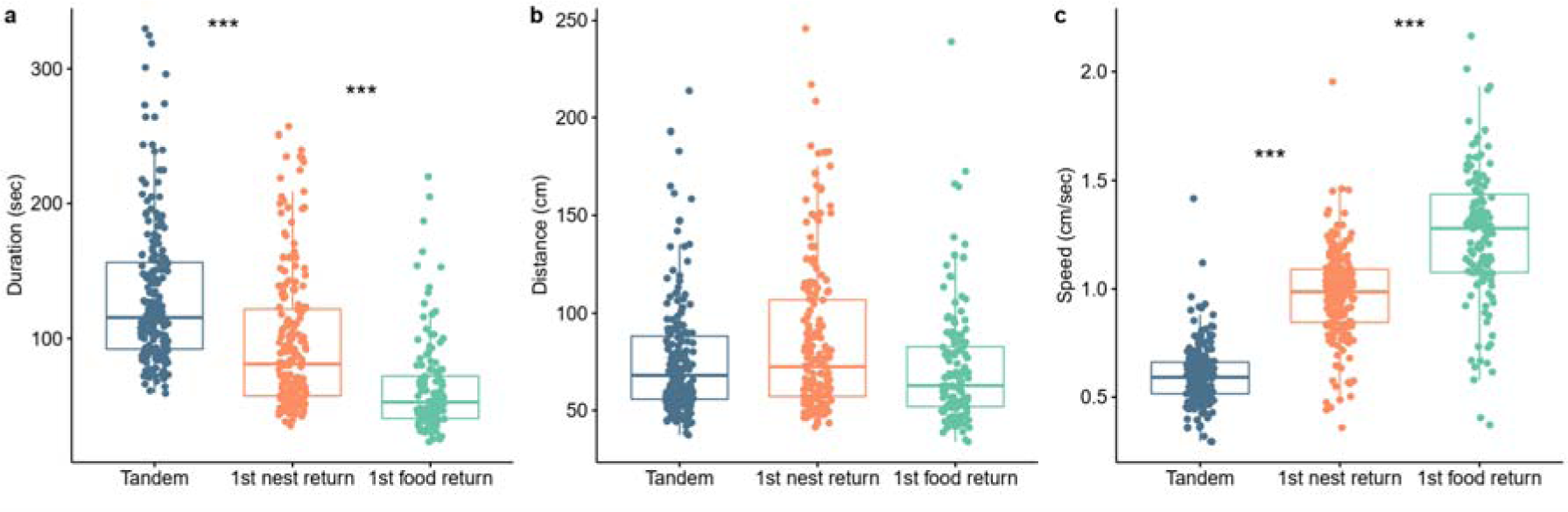
Characteristics of analysed trips during a trial. (a) The duration of trips decreased during a trial (pair-wise comparisons: to nest vs. to food, z-value = −9.69, p < 0.0001; tandem vs. to food, z-value = - 17.67, p < 0.0001; tandem vs. to nest, z-value = - 8.66, p < 0.0001). (b) The distance walked by ants did not change over time. (c) The speed of ants increased during a trial (pair-wise comparisons: to nest vs. to food, z-value = −9.57, p < 0.0001; tandem vs. to food, z-value = - 27.05, p < 0.0001; tandem vs. to nest, z-value = - 19.0, p < 0.0001).

#### Tandem run effects

Tandem pairs had no significant preference for the left or right side of the arena (44.4% of time on left side, 55.6% on right side) (binomial GLMM: z-value = −1.4, p = 0.16). The proportion of time a tandem pair spent walking on one side of the arena significantly affected the probability that ants would spend the majority of time on the same side when they returned to their nest (trip 1; Fig. 6a) (binomial GLMM: z-value = 3.33, p = 0.0009), independently of visual enrichment (z-value = 0.70, p = 0.48; interaction: LRT = 1.74, p = 0.19).

**Figure 6:**
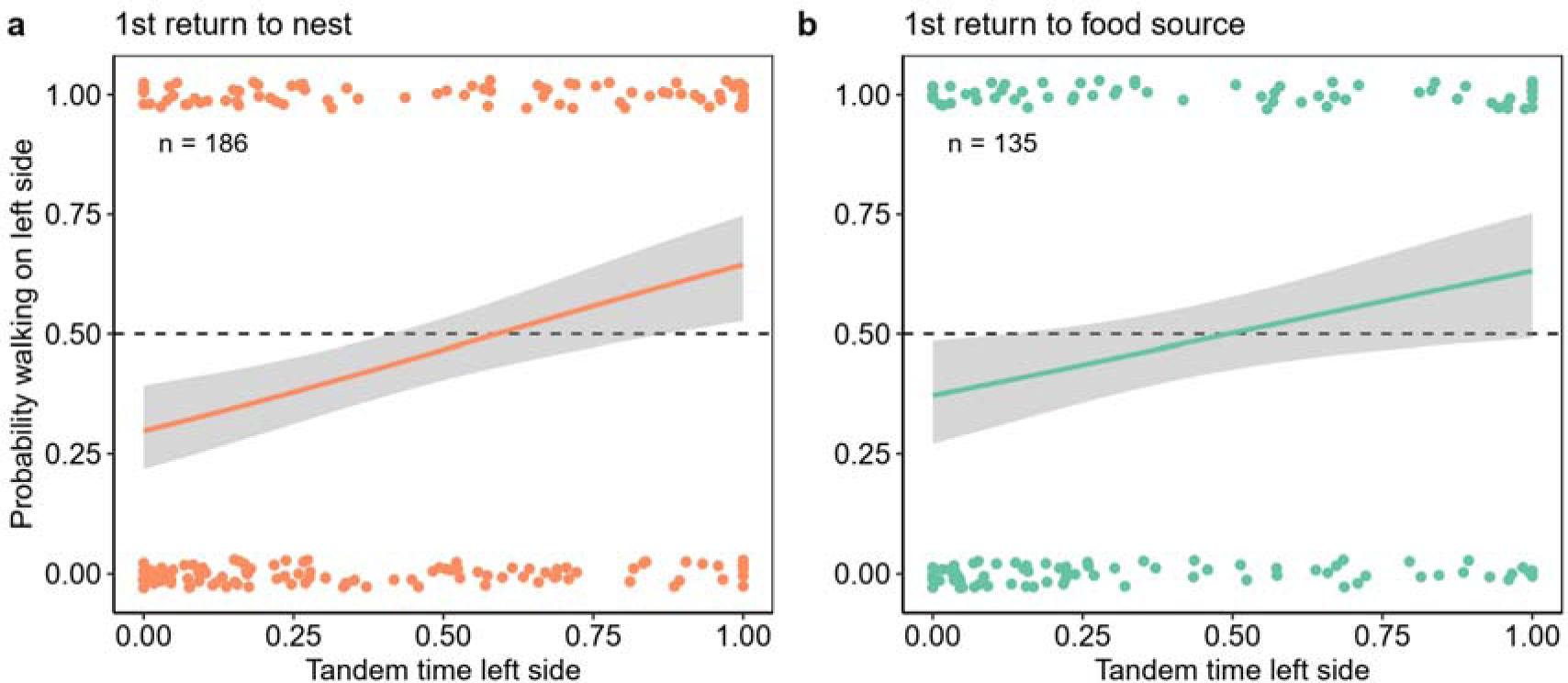
Proportion returning on left side depending on the proportion of time tandem runs spent on left side. (a) First return to the nest (trip 1) and (b) first return to the food source (trip 2) are shown. Dots represent individual ants, shaded area represents the 95%-confidence interval.

To explore whether this could be caused by chemical marks left on the ground by tandem pairs, we tested if homing ants were more likely to choose the side that was used by the previous tandem pair *(i.e.* the tandem run that happened just before the ant returned to the nest). Homing ants were not more likely to choose the side that was used by the previous tandem pair (binomial GLMM: z-value = 0.55, p = 0.58). Visualising all trajectories by ants during a typical trial further shows that colonies did not form a pheromone trail during a trial (Fig. S1; see also Sasaki et al., 2020).

We then tested if the proportion of time a tandem pair spent walking on one side also affected the probability that an ant walks on the same side when she returned to the food source alone for the first time (trip 2) and found a significant positive relationship (Fig. 6b) (binomial GLMM: z-value = 2.05, p = 0.04), independently of visual enrichment (z-value = - 0.93, p = 0.35; interaction: LRT = 1.56, p = 0.21).

We tested if the duration of the tandem run predicted the time the former follower needed to return to the nest after drinking (Trip 1) and found a significant negative relationship between tandem run duration and nest return duration (Fig. 7a) (LME: t-value = - 2.45, p = 0.015), but no effect of landmark presence (t-value = 0.84, p = 0.40; interaction: LRT = 1.39, p = 0.24). Similarly, the duration of the first return trip to the food source (trip 2) by an ant was negatively associated with tandem run duration (Fig. 7b) (LME: t-value = −2.38, p = 0.019). The presence of added visual cues significantly increased the time ants needed to return to the food source for the first time by 29% (t-value = 3.97, p = 0.0009; interaction: LRT = 1.24, p = 0.27).

**Figure 7:**
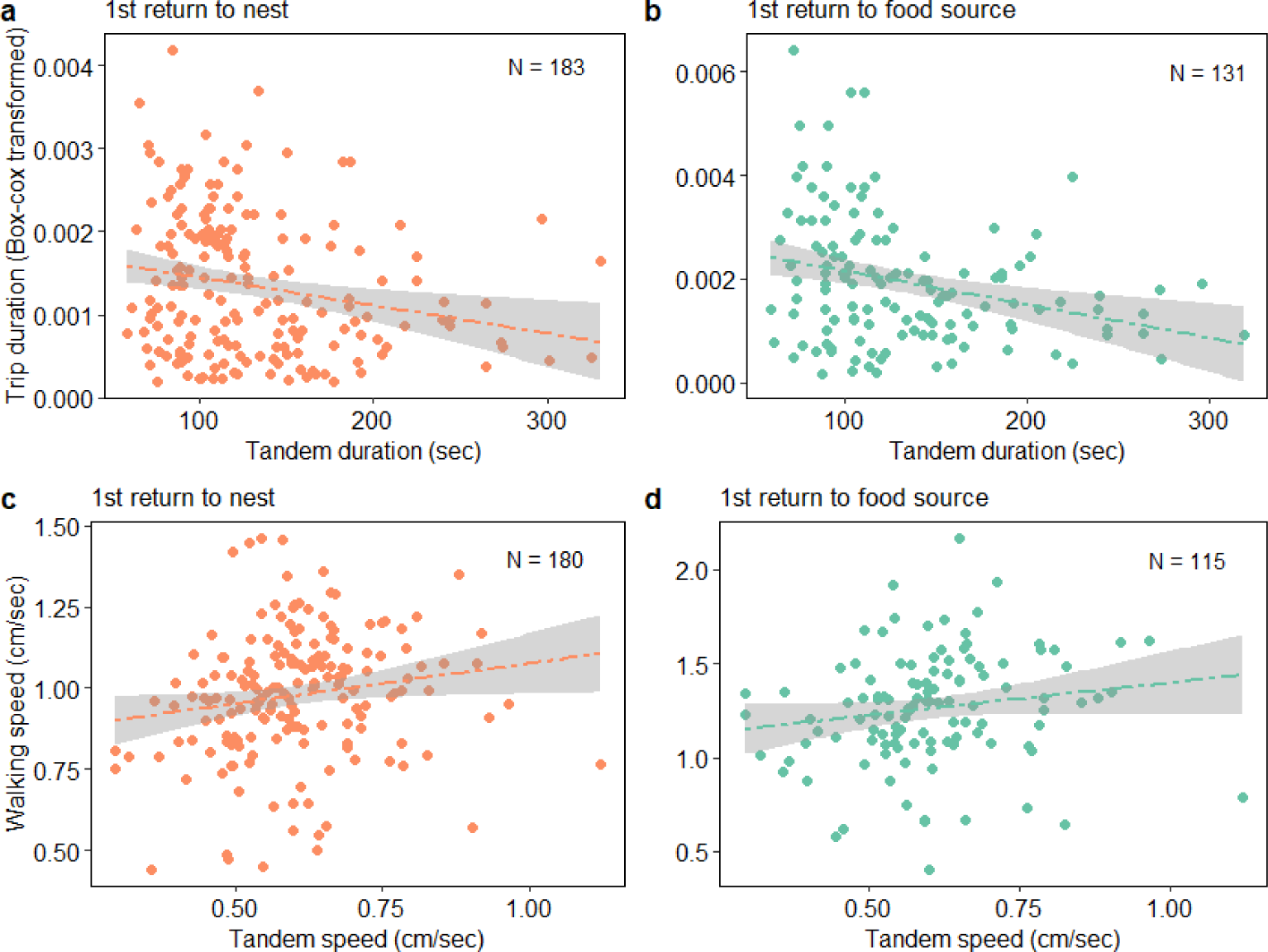
Relationship between tandem duration (sec) and speed (cm/sec) with the duration and speed of the first return trip to the nest (a,c) and the first return trip to the food source (b,d).

The distance walked by ants returning to their nest did not depend on the distance walked by the tandem pair (t-value = −1.53, p = 0.127, visual cues: t-value = 0.99, p =0.33; interaction: LRT: 0.0, p = 0.98). Likewise, the distance walked by ants returning to the food source was unaffected by the distance walked by the tandem pair (t-value = −1.38, p = 0.17), however, the presence of additional visual cues increased the walked distance by 25% (t-value = 4.32, p < 0.0001; interaction: LRT = 0.44, p = 0.51). The walking speed of former followers was weakly linked to tandem speed: Faster tandem runs predicted a faster walking speed of ants returning to their nest (Fig. 7c) (LME: t-value = 2.08, p = 0.039; visual cues: t-value = 1.18, p = 0.24; interaction: LRT = 0.96, p = 0.33) and those returning to the food source (Fig. 7d) (t-value = 1.72, p = 0.088; visual cues: t-value = 0.68, p = 0.50; interaction: LRT = 0.028, p = 0.87).

## Discussion

We found that acorn ants (*Temnothorax nylanderi*) socially learn routes during tandem running and use this information later when travelling in both directions between a food source and their nest. This contrasts with findings from another ant species, *T. albipennis,* where navigational learning during tandem running only affected routes of ants walking from the nest to the food source (Sasaki et al., 2020), but not when returning to the nest (Basari, Bruendl, et al., 2014; Franklin & Franks, 2012; Franks & Richardson, 2006; Sasaki et al., 2020).

In Experiment 1, we found that the branch taken by the tandem leader affected subsequent branch choices of their tandem followers as 67%-88% of former followers continued to use the same branch during the next eight trips (Fig. 2). Similar results were found in Experiment 2, which used an open foraging arena with a featureless floor. The arena side primarily used by the tandem pair became the preferred side of former followers on subsequent trips, both towards the nest and towards the food source (Fig. 6). This social learning seems to be remarkably efficient given that ants that discovered a food source by following a tandem run needed ∼20% less time to return to their nest than scouts that discovered the food source by themselves, *i.e.* through individual learning (Fig. 4). When former followers became tandem leaders themselves, they used the path they were taught by their tandem leaders in 90% of instances, suggesting that social learning of foraging routes spreads to the wider forager population.

These findings suggest that *T. nylanderi* foragers navigate using memorised routes, rather than path integration. Path integration is particularly useful in landscapes that do not offer many visual features, such as landscapes experienced by desert ants (Collett et al., 2013; Knaden & Graham, 2016; Müller & Wehner, 1988; Zeil, 2012, p. 201), but might be less useful when travelling on forest floors covered in physical obstacles that force ants onto tortuous paths. Retracing your steps, e.g. using idiothetic cues such as leg or body movement, might be a better strategy in such an environment. Differences in habitats could explain why route learning seems to be more dominant in acorn ants (*T. nylanderi)* than in rock ants (*T. albipennis*). The more open grasslands inhabited by *T. albipennis* (Seifert, 2018) are likely to favour a more important role of visual cues combined with path integration, especially during homing when views experienced by ants differ from those seen during the tandem run (Franklin, 2014; McLeman et al., 2002; Pratt et al., 2001; Sasaki et al., 2020). Indeed, *T. albipennis* do not follow the tandem route when returning to their nest (Sasaki et al., 2020). Studying two different *Temnothorax* species, Alleman et al. (2019) found a species specific upregulation in the expression of genes linked to learning in tandem followers, and future studies could compare brain gene expression in *T. nylanderi* and *T. albipennis* to test if different cognition-linked genes are upregulated in the brains of tandem followers and leaders in these two species.

Trip durations decreased over time in both experiments (Fig. 3, Fig. 5a). This was not due to tandem paths becoming more direct (Fig. 5b), but due to an increase in walking speed over successive trips (Fig. 5c). This is consistent with the observation that the paths of tandem leaders did not change in straightness over successive trips (Glaser & Grüter, 2018), but again contrasts findings in *T. albipennis,* where paths of tandem leaders and individual ants became more direct over successive trips (Franklin & Franks, 2012; Franks & Richardson, 2006; Sasaki et al., 2020). We speculate that a stronger reliance on target location learning and path integration in *T. albipennis* could again explain this difference between the two species. We also found that followers experiencing longer-lasting tandem runs needed less time to travel between nest and food source on subsequent trips (Fig. 7a,b). Longer tandem runs might allow followers to acquire better route information, which in turn reduces trip durations on later trips. The walking speeds of ants travelling alone between nest and food source correlated positively with the speed of the tandem run (Fig. 7c,d). Individual differences in walking speed could explain such correlations, for example caused by differences in body size, which have been shown to affect walking speed in this species (Wagner et al., 2021).

Little is currently known about the importance and identity of environmental cues for navigation in *T. nylanderi*. Adding visual cues did not affect most of the measured parameters, but we found that visual enrichment (coloured paper along the foraging arena walls in Experiment 2) caused trips back to the food source (trip 2) to be longer in both distance and duration (25% and 29%, respectively). Buehlmann et al. (2018) similarly found that desert ants walked more sinuously and slowly when encountering unexpected visual cues. In contrast, *Lasius niger* foragers travelled faster between their nest and a food source when additional visual cues were present (Grüter et al., 2015). Future research could explore whether visual enrichment initially leads to more tortuous paths, as ants need to process a larger amount of visual information, but eventually helps ants walk faster over time (see Franklin et al. 2011 for a similar effect in *T. albipennis*).

An alternative to route learning could be navigation by following pheromone trails deposited by the tandem pair. While chemical signals are known to be important during tandem running, they are short-lived and function to maintain contact between tandem partners (Basari, Laird-Hopkins, et al., 2014; Franks et al., 2022; Möglich, 1979; Möglich et al., 1974; Traniello & Hölldobler, 1984). Furthermore, chemical cues left on the ground, e.g. in the form of footprints, seem to be important for territorial marking (Bowens et al., 2013), while visual cues have been shown to dominate navigation in *T. albipennis* and *T. rugatulus* (Bowens et al., 2013; Pratt et al., 2001; Sasaki et al., 2020). We explored whether colony-specific or individual-specific (Mallon & Franks, 2000) pheromone trails could explain path choice in *T. nylanderi*. Homing ants were not more likely to choose the side that was used by the preceding tandem run *(i.e.* the tandem run that happened just before an ant left the food source to return to the nest) and the return trip distance of an ant did not correlate with the distance of her tandem run, suggesting that ants used neither colony-specific nor individual-specific pheromone trails laid by tandem pairs (see also Fig. S1).

Taken together, our data suggest that tandem followers socially learn routes and that this learning affects routes taken in both directions over several successive foraging trips. We found nuanced, but noteworthy differences between *T. nylanderi* and *T. albipennis* and speculate that these are linked to differences in the habitats used by these two species. More research is needed to understand the importance of idiothetic and environmental cues for navigation in *T. nylanderi*. We also encourage the study of navigational learning in other tandem running species, especially phylogenetically more distant species with better vision and inhabiting more varied habitats, such as *Diacamma* (Kaur et al., 2012; Mukhopadhyay et al., 2019; Mukhopadhyay & Annagiri, 2021) or *Pachycondyla* (Grüter et al., 2018; Silva et al., 2021).

## Acknowledgements

We would like to thank Jaqueline Sahm for help with the experimental set up. S.M.G. and C.G. were funded by the Deutsche Forschungsgemeinschaft (DFG) (GR 4986/1-1).

---

**Fig. S1.**
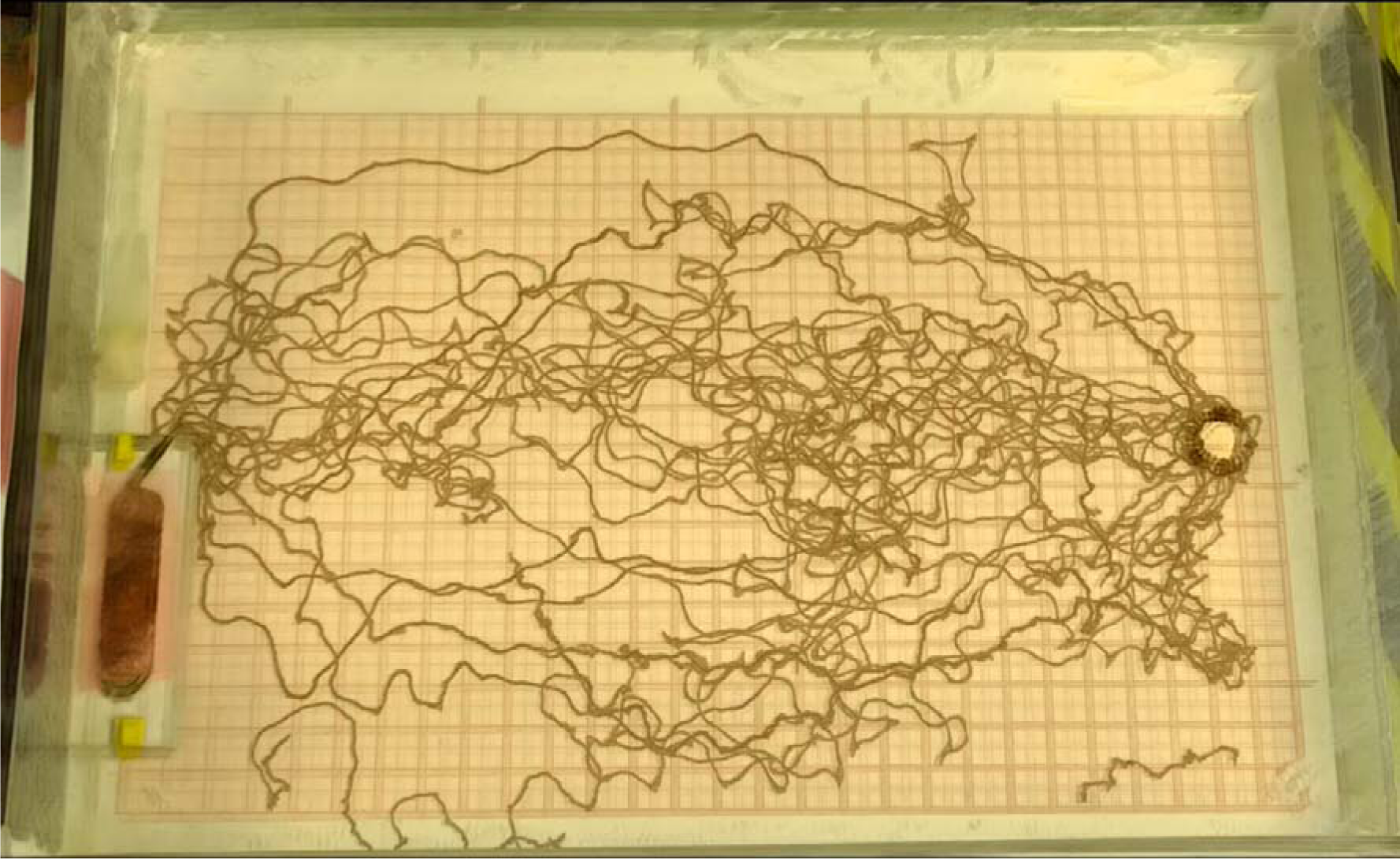
Lines show the combined paths that marked ants travelled between the nest (left side) and a sucrose solution food source (right side), after they followed a tandem run, during a 90-minute trial (Photo by Aina Colomer-Vilaplana).

